# CD36^hi^ monocytes play immunoregulatory roles in human umbilical cord blood

**DOI:** 10.1101/461459

**Authors:** Jessica G. Lee, Kathleen E. Jaeger, Yoichi Seki, Alexander J. Nelson, Alexandra Vuchkovska, Michael I. Nishimura, Paula White, Katherine L. Knight, Makio Iwashima

## Abstract

The fetal and neonatal immune systems are uniquely poised to generate tolerance to self, maternal, and environmental antigens encountered in the womb and shortly after birth. The tolerogenic nature of fetal and neonatal immunity is a rising health concern with the spread of vertically transmitted viruses, such as the Zika virus. A variety of mechanisms contribute to fetal and neonatal tolerance, including a propensity to generate Foxp3^+^ regulatory T cells (Tregs). Here, we demonstrate that a subset of CD14^+^ monocytes expressing the scavenger molecule, CD36, is able to generate CD4^+^ and CD8^+^ T cells that express Foxp3 from umbilical cord blood (UCB). Monocyte-induced Foxp3^+^ T cells have potent suppressive functions on T cell proliferation and maintain Foxp3 expression over six weeks *in vitro*. Importantly, UCB-derived Foxp3^+^ T cells are distinguishable from adult peripheral blood (APB) CD4^+^CD25^+^ Tregs by surface antigen expression. While UCB-derived Foxp3^+^ T cells express prototypic Treg-associated surface antigens, such as CD25 and glucocorticoid-induced tumor necrosis factor-related receptor (GITR), only UCB-derived Foxp3^+^ T cells express CD26. In addition, most UCB-derived CD8^+^Foxp3^+^ T cells express CD31. Mechanistically, both APB and UCB-derived monocytes support the development of Foxp3^+^ T cells from naïve T cells, but APB naïve T cells are less efficient in expressing Foxp3 than UCB naïve T cells. These data suggest that antigen presentation by CD36^hi^ monocytes in the fetus leads to the development of a group of T cells that share some but not all phenotypes of adult thymus-derived Tregs.

## 1. Introduction

The concept of fetal tolerance was introduced in 1945 when R. D. Owen noted dizygotic cattle twins can share a placenta and maintain blood cells from their sibling [1]. Each adult cow subsequently tolerates blood cells from its genetically disparate twin. Similarly, Medawar and Billingham demonstrated that mice and chickens that receive allogeneic tissues *in utero* are tolerant to the donor as adults, but maintain the ability to reject skin grafts to which they have not been tolerized [2]. These findings suggested fetuses induce antigen-specific immune tolerance to antigens encountered *in utero* and this tolerance can be maintained into adulthood.

Multiple mechanisms contribute to fetal tolerance, including clonal deletion, anergy, differences in APCs, and Foxp3^+^ regulatory T cells (Tregs) generation (reviewed by [3][4]). Tregs limit immune responses against self and non-self antigens and are essential for immune homeostasis. Tregs can develop in the thymus (thymus derived Tregs, tTregs) or can be induced from peripheral naïve CD4^+^ T cells *in vivo* (peripheral Tregs; pTregs) or *in vitro* (induced Tregs; iTregs) [5]. Human umbilical cord blood (UCB) can be a source of iTregs and fetal CD4^+^ T cells have a strong tendency to differentiate into regulatory cells [6-8]. This is likely due to both T cell intrinsic factors and signals from UCB APCs [9, 10]. However, the mechanisms leading to robust iTreg generation from the fetus are not well understood.

Here, we demonstrate human UCB contains a subset of monocytes (CD14^+^CD36^hi^) that is highly tolerogenic and induces stable CD4^+^ and CD8^+^ Foxp3^+^ T cells with regulatory functions. CD14^+^CD36^hi^ monocytes express membrane-bound TGF-β and can metabolize vitamin A into retinoic acid, a known inducer of Tregs. Adult peripheral blood (APB) derived monocytes are equally capable of inducing Foxp3^+^ T cells but adult T cells are less effective in becoming Foxp3^+^ T cells. Notably, while Foxp3^+^ T cells from UCB are potent in suppressing conventional T cell proliferation, they differ from adult peripheral blood Foxp3^+^ Tregs in surface antigen expression. These data suggest that fetal peripheral blood is uniquely conditioned to generate immunoregulatory adaptive responses against incoming antigens by promoting a unique group of T cells in a monocyte-dependent manner.

## 2. Materials and Methods

### 2.1 Antibodies

Antibodies used for flow cytometry were: anti-phospho-SMAD2/3 (BD Biosciences; San Jose, CA), anti-IL-2 (BD Biosciences), anti-LTBP1 antibody (R&D Systems; Minneapolis, MN), anti-CD127 (eBioscience; San Diego, CA). Anti-CD4, CD8, CD14, CD16, CD25, CD26, CD31, CD36, CD64, OX-40, GITR, LAP, TNF, GM-CSF, IL-8, and IFN-γ antibodies were from BioLegend (San Diego, CA). Anti-Foxp3 was from both BD Biosciences and BioLegend. Functional grade antibodies for cell culture, anti-CD3 (OKT3) and anti-CD28 were from BioLegend.

### 2.2 Chemicals and recombinant proteins

The following reagents were used: recombinant human IL-2 (PeproTech; Rocky Hill, NJ); a TGF-β receptor I kinase inhibitor, SB431542 (Sigma-Aldrich; St. Louis, MO); an RAR antagonist, LE135 (Tocris; Bristol, UK); and a RAR agonist AM580 (Tocris). Carboxyfluorescein succinimidyl ester (CFSE) was purchased from Invitrogen (Carlsbad, CA).

### 2.3 cTreg Induction Culture

Total UCB mononuclear cells were stimulated with human IL-2 (10 ng/ml; >100 U/ml) and anti-CD3 (0.2 µg/ml) in RPMI 1640 (GE Healthcare Hyclone, Chicago, IL) supplemented with 10% fetal calf serum and essential- and non-essential amino acids. Cells were split with a media change every 2-3 days, maintaining IL-2 concentrations. cTreg percentages were analyzed after 12-15 days of culture, unless otherwise specified. For purified co-cultures, T cells were cultured with monocytes at a 1:3 ratio with anti-CD3 and IL-2. For bead-based stimulation, T cells were cultured with polystyrene beads (ration: 4 beads to 1 T cell) pre-coated with anti-CD3 and anti-CD28 Abs (10µg/ml each). Where indicated, SB431542, LE135, or AM580 were added at the listed concentrations at the beginning of cultures, unless otherwise specified.

### 2.4 Mononuclear Cell Isolation and Cell purification

UCB was collected into citrate phosphate dextrose solution. Adult PBMCs from healthy donors were collected in heparin or buffered sodium citrate. Mononuclear cells were enriched by density dependent centrifugation using Lymphocyte Separation Medium (Corning Cellgro, Tewksbury, MA). In some samples, RBCs were lysed with ACK lysis buffer (Gibco, NY). For UCB, a second isolation with Lymphocyte Separation Medium was performed after ACK lysis. Total CD3^+^ T cells, CD4^+^ T cells, naïve CD4^+^ T cells and CD14^+^ monocytes were enriched from mononuclear cells using BD IMag Enrichment Sets (BD Biosciences) or EasySep enrichment kits (STEMCELL Technologies; Vancouver, Canada). IMag kits were used for depletion of CD14^+^ and CD3^+^ cells (for Cell Titer Glo assay, Promega). CD3 depleted (for CFSE assay), CD14^+^CD36^hi^, and CD14^+^CD36^lo^ cells were isolated by cell sorting on FACS Aria (BD Biosciences). EasySep Human CD25 depletion kit was used for depletion of CD25^+^ cells (STEMCELL). Adult Tregs were enriched using EasySep Human CD4^+^CD127^low^CD25^+^ Regulatory T cell kit (STEMCELL).

### 2.5 Flow Cytometry

Foxp3 staining was performed with the Foxp3 Fix/Perm Buffer Set (Biolegend). For phospho-Smad2/3 staining, cells were fixed using Lyse/Fix Buffer (BD Biosciences) and permeabilized with Perm Buffer III (BD Biosciences). Surface stains were performed using standard protocols. Data were collected on a FACS Canto II (BD Biosciences) or FACS LSRFortessa (BD Biosciences) and analyzed using FlowJo software (Tree Star, Inc., Ashland, OR).

### 2.6 Cytokine Assay

Cells were restimulated for 4 hours using phorbol myristate acetate (PMA; 50 ng/ml; Sigma Aldrich) and ionomycin (1 µM; Sigma Aldrich) in the presence of monensin (1 µM; Biolegend) in RPMI 1640 (Hyclone) supplemented with 10% fetal calf serum. Cells were fixed and permeabilized using Foxp3 Fix/Perm Buffer Set (Biolegend) followed by intracellular staining for Foxp3 and cytokines according to manufacturer instructions.

### 2.7 TGF-β Bioassay

MFB-F11 cells (TGFβ-1 deficient MEFs transfected with a SMAD-binding element promoter fused to secreted embryonic alkaline phosphatase) were described previously (provided by Dr. Tesseur, Stanford University) [11] and used as reported.

### 2.8 Suppression Assay (cell division)

Treg suppressive function was tested by 2 methods: CFSE dilution and Cell Titer Glo® assay. For the CFSE dilution assay, Tregs were generated by UCB stimulation with anti-CD3 and IL-2, as described above. Foxp3 expression was confirmed by flow cytometry and CD4^+^ and CD8^+^ cells were separated by immunomagnetic enrichment. Unstimulated naïve CD4^+^ T cells were enriched from allogeneic adult PBMCs, labeled with 5µM CFSE and used as responder cells. CD3 depleted cells sorted from the same PBMC donor and irradiated at 3000 rad were used as APCs. Responder cells were stimulated with anti-CD3 (0.2 µg/ml) and APCs at a 1:1 ratio, in the presence or absence of the indicated ratio of Tregs. The percent of proliferating cells was determined after 5 days of culture as the percent of cells with diluted CFSE using flow cytometry. For the Cell Titer Glo assay, unstimulated CD4^+^ T cells enriched from the same donor were used as responder cells. Irradiated syngeneic T cell depleted samples were used as APCs. Cell proliferation was determined after 7-10 days of culture, using the Cell Titer Glo® Luminescent Cell Viability Assay (Promega, Madison, WI), according to the manufacturer’s protocol.

### 2.9 Suppression assay (Cell growth)

To perform a proliferation assay, CD4^+^ CD25^+^ Tregs were enriched from indicated day 14 cultures via the Human Regulatory T Lymphocyte Separation Set – DM (BD IMag). Additionally, total CD4^+^ T cells were enriched as previously described from syngenic samples that had been frozen since the day of collection. Cells were cultured in 96 well U-bottom plates at increasing Treg:Responder T cell ratios in cell culture media with soluble anti-CD3 (Biolegend) at 0.2 ug/ml. The same number of responder CD4^+^ T cells were cultured in each well with an increasing ratio of Tregs added to each subsequent well. Wells of either responder CD4^+^ T cells or Tregs alone were cultured as controls for proliferation. Culture was maintained for 7-10 days at 37°C until measure of cell proliferation by Cell Titer Glo® Luminescent Cell Viability Assay (Promega) according to protocol.

### 2.10 ALDEFLUOR™ Assay

ALDH activity was measured in freshly isolated UCB mononuclear cells using the ALDEFLUOR™ Kit (STEMCELL Technologies), according to the manufacturer’s instructions.

### 2.11. Statistical Analysis

The majority of the statistical analyses were performed using GraphPad Prism software with one way ANOVA (San Diego, CA). The following designation was used throughout the paper: * p <0.05, ** p < 0.01, *** p < 0.001, **** p < 0.0001, unless otherwise specified. Where indicated, the percent inhibition of Treg differentiation was calculated as [(%Treg control - %Treg treated)/ %Treg control].

## 3. Results

### 3.1 Induction of Foxp3^+^ Tregs from human UCB

To understand the cellular mechanisms that establish fetal tolerance, we first determined whether full term fetuses have more Foxp3^+^ CD4^+^ cells than adults. The frequency of CD4^+^ Foxp3^+^ cells was similar in UCB and adult APB (Fig. 1A). However, when total mononuclear cells were stimulated with an anti-CD3 antibody in medium containing IL-2 over the course of two weeks, a large percentage of UCB CD4^+^ T cells (65.6%, ±3.5) acquired the prototypic Treg markers CD25 and Foxp3, whereas few (19.5%, ± 7.3) adult CD4^+^ cells became Foxp3^+^ (Fig. 1B, C). Notably, stimulated UCB also gave rise to CD8^+^ Foxp3^+^ T cells (Fig. 1B, C). CD8^+^ suppressive cells have been previously described, however not derived from UCB cells [12]. Foxp3^+^ CD4^+^ and CD8^+^ T cells induced from UCB were not derived from pre-existing Tregs, as removal of CD4^+^ CD25^+^ T cells did not reduce the frequency of Foxp3^+^ T cells after anti-CD3 stimulation of UCB (Fig. 1D, E).

**Fig. 1.**
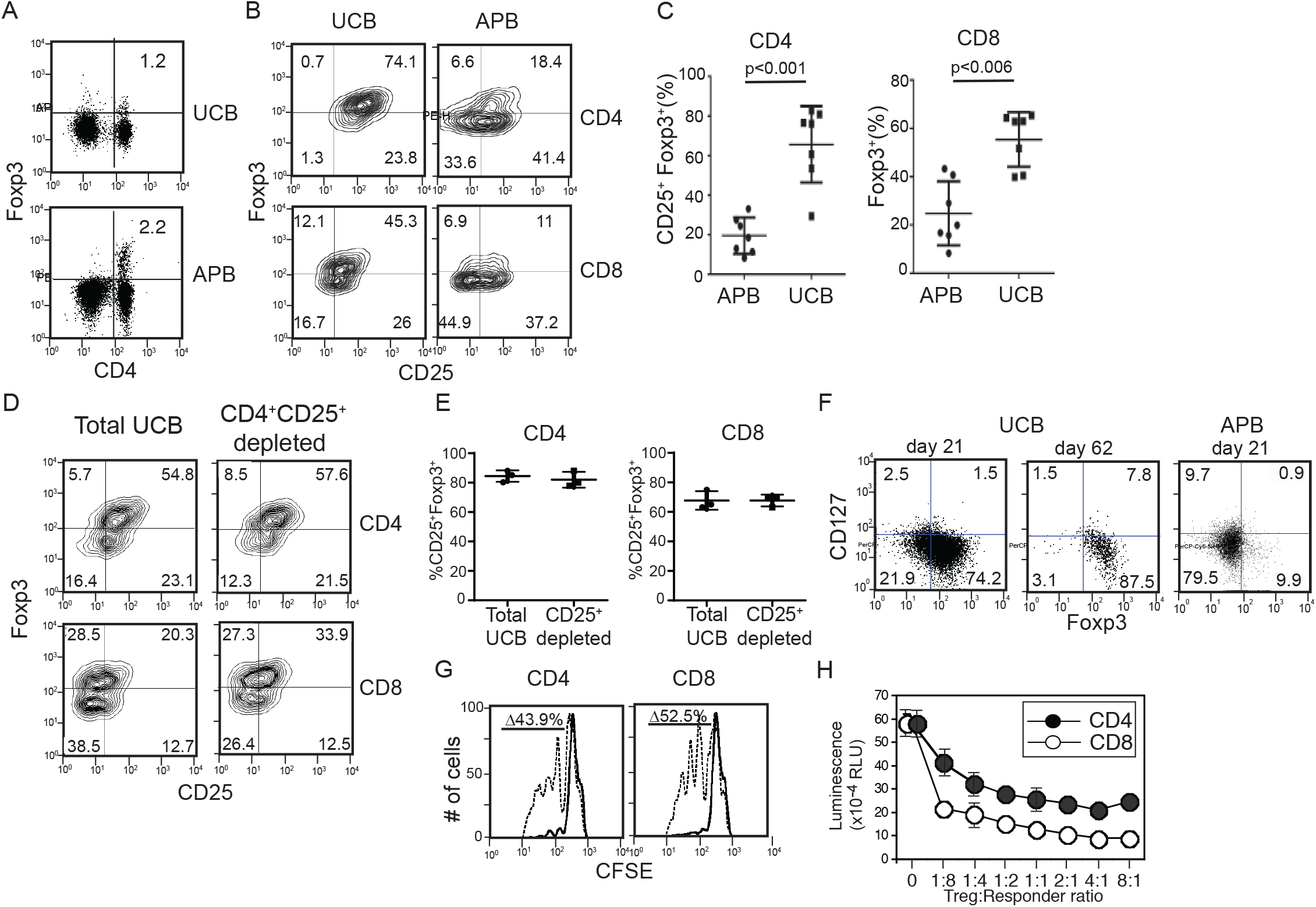
Induction of immunoregulatory Foxp3^+^ T cells from UCB. (**A**) Foxp3 expression among freshly isolated UCB and adult APB. **(B-C)** Frequency of CD4^+^ and CD8^+^ cells expressing Foxp3 after anti-CD3 and IL-2 stimulation and expansion for 14 days of APB or UCB, n=7 each; 2-tailed Student *t* test. (**D-E**) Frequency of cells expressing Foxp3 and CD25 from stimulation of total UCB mononuclear cells and cells depleted of CD25^+^ cells prior to stimulation. (**F**) Stability of Foxp3^+^ CD127-phenotype on CD4^+^ cells from UCB or APB. T cells stimulated as in (B) were expanded in the medium containing IL-2 and were analyzed 21 or 62 days after stimulation. Adult blood was not analyzed at day 62, since the majority cells did not express Foxp3 at day 21. (**G-H**) *In vitro* suppression of T cell proliferation by UCB derived CD4^+^ and CD8^+^ Foxp3^+^ T cells. UCB-derived Foxp3^+^ T cells were induced as in (B) and used in a standard suppression assay. CFSE-labeled, allogeneic adult naïve CD4^+^ T cells were used as responder cells. Responder to Treg ratio was 1:1. (G) Representative CFSE histograms of gated responder cells in the presence or absence of CD4^+^ or CD8^+^ T cells from UCB. (H) Summarized data from 3 UCB donors, one-way ANOVA with Dunnett’s multiple comparisons test.

In contrast to murine T cells, Foxp3 and CD25 are transiently expressed on activated human CD4^+^ T cells, with only Tregs stably expressing Foxp3. Therefore, we tested the stability of Foxp3 expression on stimulated UCB T cells by extending the culture with IL-2 beyond two weeks. To differentiate Foxp3^+^ T cells from effector T cells, we concurrently assessed the presence of the IL-7 receptor *α* chain (CD127), which is expressed by conventional CD4^+^ T cells but not Tregs [13]. Activated CD4^+^ UCB cells sustained the Foxp3^+^ CD127-phenotype after both 21 and 62 days in culture, while activated CD4^+^ cells from APB had few Foxp3^+^ CD127-cells after only 21 days of culture (Fig. 1F). These data demonstrate stimulated UCB CD4^+^ T cells maintain a Treg-like phenotype for more than two months in culture.

To determine if the Foxp3^+^ cells induced from UCB have a regulatory function, we examined their suppressive activity by *in vitro* suppression assays (Fig. 1G, H). Naïve CD4^+^ responder T cells proliferate vigorously in response to anti-CD3 antibody stimulation; however, the addition of Foxp3 expressing CD4^+^ or CD8^+^ T cells from anti-CD3 and IL-2 stimulated UCB cells substantially reduced cell proliferation. Suppression was observed when the ratio of stimulated UCB T cells (Treg) to naïve T cells (responder) was as low as 1:8 (Fig. 1H), showing the Foxp3^+^ cells generated from UCB are functional regulatory cells. While activated T cells are known to have suppressive functions on T cell proliferation, Treg-depleted pre-activated PBMC T cells did not reduce responder T cell proliferation under the same conditions, showing that T cell suppression by Foxp3^+^ UCB T cells is not caused because they are activated T cells (sFig. 1).

### 3.2 UCB Derived Foxp3^+^ T cells uniquely express CD26 and CD31

To compare UCB derived Foxp3^+^ T cells and adult Tregs, we assessed surface antigen expression. We have recently determined that UCB naïve T cells express Dipeptidyl Peptidase 4 (CD26) while adult naïve T cells do not [14]. Similarly, UCB Foxp3^+^ T cells maintained expression of CD26 (Fig. 2A). Practically all UCB Foxp3 expressing CD4^+^ and CD8^+^ T cells expressed CD26 (100%, 99.9%±0.03, respectively; Fig. 2A, C). In contrast, CD26 was not detectable on the majority of adult Tregs (Fig. 2A, C). CD26 is expressed at high level by several T cell subsets including Th17 cells [15] and mucosal-associated invariant T cells (MAIT cells) [16]. Gene knockout mouse studies indicate that CD26 plays both T cell activating [17] and modulatory functions [18].

**Fig. 2.**
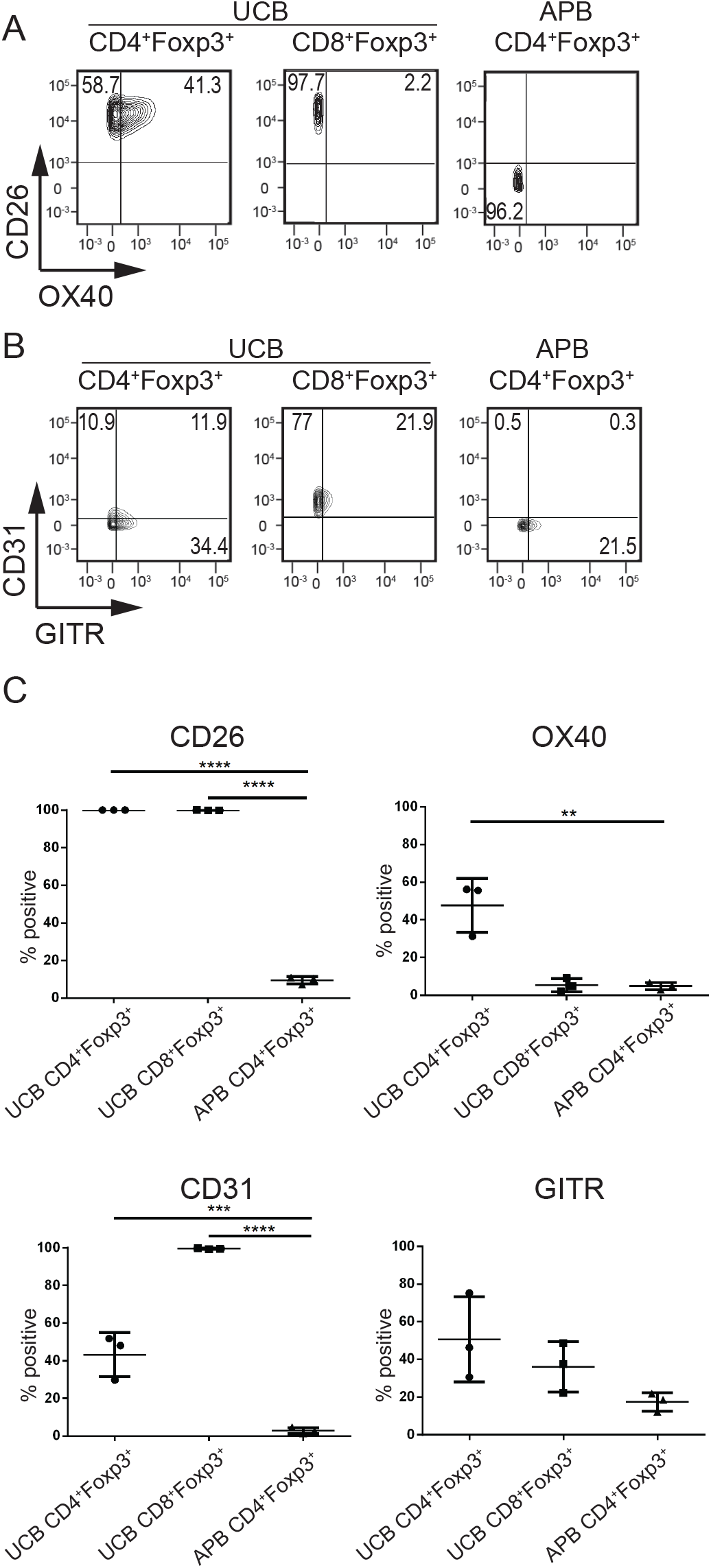
Surface antigen expression by Foxp3^+^ T cells derived from UCB. (**A, B**) Representative data from surface antigen expression of CD26, OX40, CD31, and GITR by UCB Foxp3^+^ T cells induced by stimulation with anti-CD3 and IL-2 and expanded for 14 days, or by freshly isolated APB Tregs. (**C**) Summarized data from (A-B), each dot represents one donor. Analyzed by one-way ANOVA with Dunnett’s multiple comparisons test.

OX40 is a co-stimulatory molecule of the TNFR family expressed on activated effector T cells and Foxp3^+^ Tregs and reportedly promotes proliferation and survival in effector T cells, but blocks the activity and differentiation of Tregs [19, 20]. A subset of UCB CD4^+^ Foxp3^+^ T cells (47.6%±8.2), but not CD8^+^ Foxp3^+^ T cells, expressed OX40 (Fig. 2A, C).

In addition, almost all CD8^+^ Foxp3^+^ T cells (99.5%±0.15) notably expressed an adhesion molecule CD31, a marker of recent thymic emigrant [21] and a regulatory molecule of T cell activation [22]. A portion of CD4^+^ Foxp3^+^ T cells (43.2%±0.15) also expressed CD31 (Fig. 2B, C). In contrast, neither OX40 nor CD31 were substantially detected on adult tTregs. Although UCB-derived Foxp3^+^ T cells can be distinguished from APB Tregs by surface antigens, they share some phenotypic similarities, including GITR expression (Fig. 2B, C), a molecule known to be upregulated by Foxp3 [23]. Based on these data, we termed cord blood-derived Foxp3^+^ T cells as cTregs (cord blood-derived Tregs) herein.

### 3.3 Role of CD14^+^CD36^+^ monocytes in cTreg induction

The high frequency of stimulated UCB T cells expressing Foxp3^+^ may be due to an intrinsic propensity of UCB T cells to differentiate into Foxp3^+^ T cells. Alternatively, other cells within the UCB mononuclear cell fraction may promote Foxp3^+^ T cell differentiation. To test whether UCB T cells are intrinsically programmed to differentiate into Foxp3^+^ T cells, we stimulated them with anti-CD3 and anti-CD28 antibody-coated beads and found the frequency of Foxp3^+^ T cells was markedly reduced compared to T cells stimulated in the presence of other UCB cells (Fig. 3A). These data show that UCB T cells are not pre-programed to be Foxp3^+^ T cells and indicate that a non-T cell population is required for efficient Foxp3^+^ T cell generation from UCB. To identify which subset of UCB cells induces Foxp3^+^ T cell differentiation, we depleted non-T cell subsets from UCB and found that CD14^+^ cell depletion significantly reduced the generation of Foxp3^+^ cells (Fig. 3B, C).

**Fig. 3.**
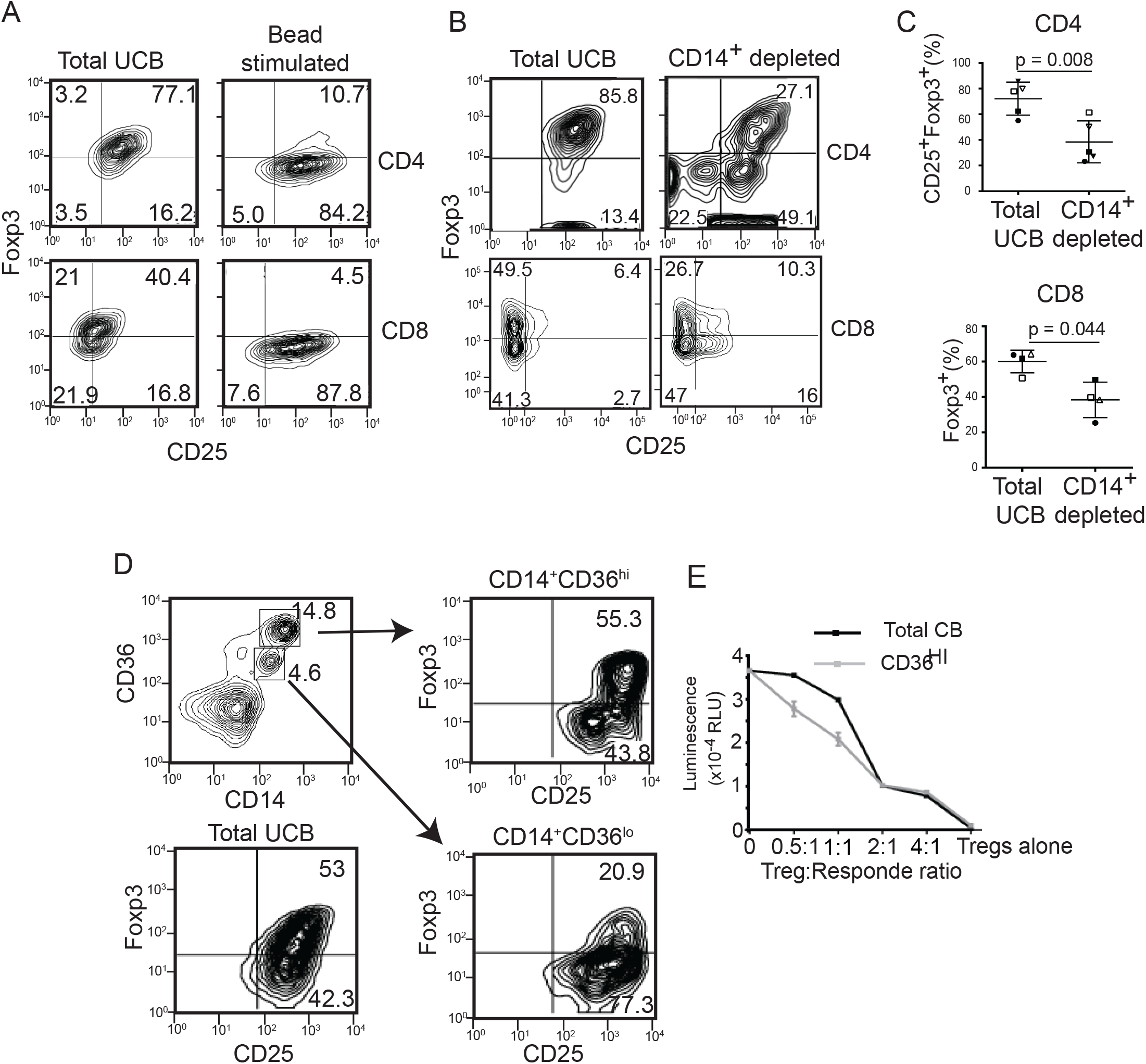
CD14^+^ cells induce Foxp3^+^ cTregs. (**A**) Foxp3 expression by purified UCB T cells stimulated with anti-CD3/anti-CD28 antibody-coated beads in the presence of IL-2 after 14 days of expansion. As a positive control, total UCB mononuclear cells from the same donor were stimulated by anti-CD3 antibody and IL-2. **(B-C)** The expression of Foxp3 on gated CD4^+^ or CD8^+^ T cells after 14 days of expansion from total UCB or CD14-depleted UCB stimulated with anti-CD3 and IL-2. **(B)** A representative plot and **(C)** summarized data using the paired Student t test. Each shape represents a different donor, n=4-5. (**D**) Expression of CD14 and CD36 among freshly isolated total UCB mononuclear cells (upper left panel). Expression of CD25 and Foxp3 by gated CD4^+^ T cells from total UCB cultures (lower left panel) or UCB naïve CD4^+^ T cells co-cultured with purified CD14^+^CD36^hi^ (upper right panel) or CD14^+^CD36^lo^ cells (lower right panel). All cultures were stimulated with anti-CD3 and IL-2 and expanded for 14 days. Data represents at least 3 independent experiments. (**E**) Suppression of T cell proliferation by T cells derived from total UCB (black line) or from CD4^+^ T cells co-cultured with CD14^+^CD36^hi^ cells (gray line), measured using the Cell Titer Glo assay.

Human monocytes are a heterogeneous population of cells. To determine the subsets of monocytes present in the perinatal blood, we examined several known surface markers and found the expression of CD36 distinguishes CD14^+^ cells into two groups: CD36^hi^ and CD36^lo^ (Fig. 3D). CD36 is a surface scavenger protein conserved in both non-vertebrate and vertebrate immune cells [24]. We next sorted UCB CD14^+^ cells by CD36 expression and determined that CD14^+^CD36^hi^ cells induce Foxp3^+^ T cells from naïve T cells at a level comparable to total UCB, while CD14^+^CD36^lo^ cells induce substantially less Foxp3^+^ cells (Fig. 3D). In contrast, CD14-CD36^+^ and CD14-CD36-cells induced negligible Foxp3^+^ cells (data not shown). Furthermore, T cells induced from co-culture with CD14^+^CD36^hi^ monocytes suppressed syngeneic T cell proliferation comparable to T cells induced from total UCB mononuclear cells (Fig. 3E). These data show CD14^+^CD36^hi^ cells are sufficient to induce cTregs. Other surface antigens often used for monocyte analysis (CD64 and CD16) [25] showed the majority of CD14^+^CD36^hi^ monocytes express CD64 and low levels or no CD16, consistent with the classical monocyte phenotype (sFig. 2).

Monocytes have not been widely recognized as cells that induce stable Foxp3^+^ T cells and therefore the mechanism was unknown. Thus, we determined whether CD14^+^CD36^hi^ monocytes use mechanisms similar to other Treg-inducing cells. TGF-β plays a critical role in iTreg induction in both human and murine models [26], however CD14^+^CD36^hi^ monocytes do not require exogenous TGF-βfor Foxp3^+^ T cell induction. To test if endogenous TGF-β is required for cTreg differentiation, we added a TGF-β receptor kinase inhibitor (SB541542) to total UCB cells stimulated by anti-CD3 and IL-2. The inhibitor blocked CD25^+^Foxp3^+^ T cell differentiation in a dose-dependent manner (Fig. 4A). Similarly, an anti-TGF-β Ab blocked cTreg differentiation (sFig. 3). To determine when TGF-β signaling is required for cTreg differentiation, we added the TGF-β receptor kinase inhibitor at various times after the beginning of T cell stimulation. Addition of the inhibitor as late as 24 hours after the start of culture blocked cTreg differentiation at a comparable level of inhibition from the start of the culture. In contrast, inhibition of TGF-β signaling 48 hours after start of culture did not reduce cTreg differentiation, suggesting that TGF-β is required for cTreg differentiation mainly between 24 and 48 hours after start of TCR stimulation (Fig. 4B).

**Fig. 4.**
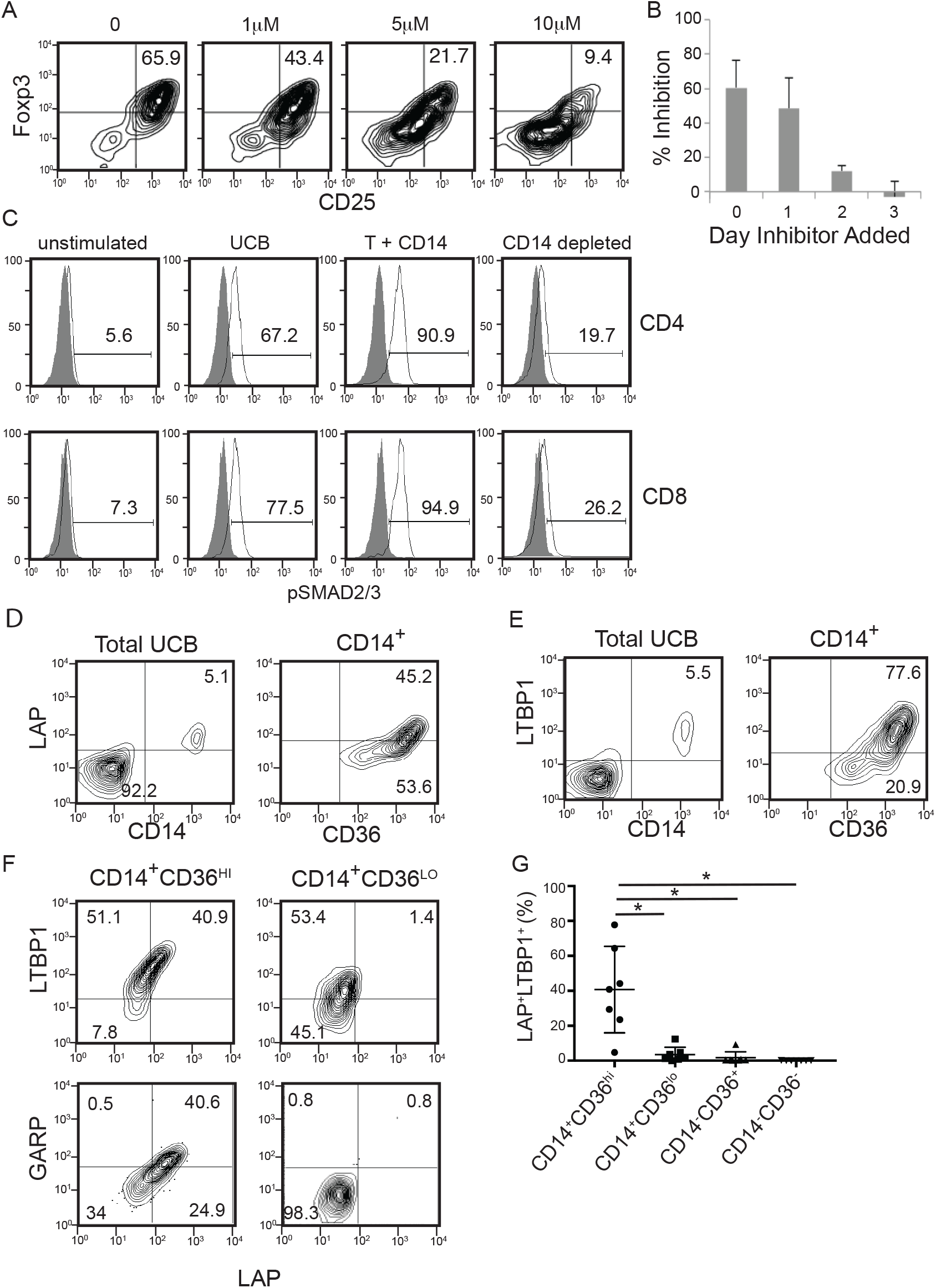
Requirement of TGF-β signaling for Foxp3^+^ cTregs induction from UCB cells. (**A-B**) Inhibition of Foxp3^+^ cTregs induction by SB431542, an inhibitor of the TGF-β receptor I kinase. (**A**) Expression of CD25 and Foxp3 by UCB CD4^+^ T cells 21 days after stimulation by anti-CD3 and IL-2 in the presence or absence of SB431542 at the indicated concentrations. (**B**) % inhibition of Foxp3^+^ CD4^+^ cTregs cell generation by SB431542, added at various times after the start of UCB stimulation with anti-CD3 and IL-2. Percent inhibition is calculated as [(%Tregs _DMSO treated_) – (% Treg _SB431542 treated_)]/ (%Tregs _DMSO treated_). Data included from 3 donors. (**C**) Activation of Smad2/3 by CD14^+^ cells. Total UCB cells, purified CD4^+^ and CD14^+^ cells from UCB, or CD14^+^ depleted UCB cells were stimulated by an anti-CD3 antibody and IL-2 or left unstimulated for 1 day. After stimulation, cells were harvested, permeabilized, and stained with antibodies against pSmad2/3. Isotype control staining is shown by the shaded areas. Data for CD4^+^ (upper panel) and CD8^+^ T cells (lower panel) are shown. Data representative of 2-3 independent experiments. **(D, E)** LAP **(D)** and LTBP1 **(E)** expression by total UCB mononuclear cells and CD14^+^ gated monocytes. (**F**) Co-expression of LAP and LTBP1/GARP by CD14^+^CD36^hi^ and CD14^+^CD36^lo^ gated cells. **(G)** Summary graph of LAP and LTBP1 co-expression by CD14^+^CD36^+^ gated subsets, one-way ANOVA with Dunnett’s multiple comparison’s test, n=7.

With this knowledge, we next determined whether monocytes are the source of TGF-β for cTregs differentiation. Activation of the TGF-β receptor leads to Smad2 and Smad3 phosphorylation [27]. We predicted that if monocytes are the endogenous source of TGF-β for Foxp3^+^ T cell differentiation, then depleting monocytes would abrogate Smad2/3 phosphorylation (pSmad2/3) in stimulated T cells. When total UCB cells were stimulated for 1 day with anti-CD3 and IL-2, we observed substantial phosphorylation of Smad2/3 in CD4^+^ and CD8^+^ T cells (Fig. 4C). Smad2/3 was also phosphorylated in T cells cultured with enriched CD14^+^ monocytes, but greatly reduced in T cells from CD14 depleted UCB cultures. These data demonstrate that CD14^+^ cells are sufficient and required for activation of Smad2/3 in UCB T cells.

TGF-β is produced in a latent form which requires activation to elicit downstream signaling [28]. We hypothesized CD14^+^CD36^hi^ monocytes secrete active TGF-βto induce Foxp3^+^ T cells. However, when we analyzed the culture supernatant of CD14^+^CD36^hi^ monocytes, active TGF-β was below the detection limit of the bioassay (∼1pg/ml) and only a limited amount of the inactive form of TGF-β (6pg/ml) was detected [11] (sFig. 4). Thus, we hypothesized CD14^+^CD36^hi^ cells present TGF-β to T cells via direct cell-to-cell contact and/or by vesicular forms that are not detectable from the culture supernatant.

To test this hypothesis, we determined the level of the TGF-β complex on the surface of CD14^+^CD36^hi^ monocytes. TGF-β is initially produced as a precursor that is cleaved into latency associated peptide (LAP) and the growth factor domain. The two peptides associate and covalently bind with Latent TGF-β Binding Proteins (LTBPs). The LTBP/LAP complex can be secreted from the cell and attached to the extracellular matrix through the interaction of LTBPs with other proteins. As predicted, CD14^+^CD36^hi^ monocytes were the major group of UCB cells that expressed LAP along with LTBP1on the cell surface (Fig. 4D, E, F). In addition, LAP^+^ CD14^+^CD36^hi^ also expressed GARP, another surface molecule known to capture the latent form of TGF-β [29] (Fig. 4F). The co-expression of LAP and LTBP-1 was significantly higher on CD14^+^CD36^hi^ monocytes than on CD14^+^CD36^lo^ monocytes or other UCB cells (Fig. 4G). All LAP^+^ cells co-expressed LTBP1 and GARP, indicating CD14^+^CD36^hi^ monocytes present LAP on the cell surface as a complex with LTBP1 and/or GARP.

### 3.4 CD14^+^CD36^hi^ monocytes express active ALDH

Previous reports showed retinoic acid, a vitamin A metabolite, supports Treg differentiation in a TGF-β dependent manner [30]. To determine whether retinoic acid is also required for UCB Foxp3^+^ T cell differentiation, we cultured UCB in the presence of a retinoic acid receptor (RAR) antagonist, LE135. The antagonist decreased Foxp3 expression in CD4^+^ and CD8^+^ T cells (Fig. 5A, B). This effect was specific to the inhibition of RAR since Foxp3^+^ T cell differentiation was restored when an RAR agonist (AM580) was added in addition to the antagonist, in a dose-dependent manner.

**Fig. 5.**
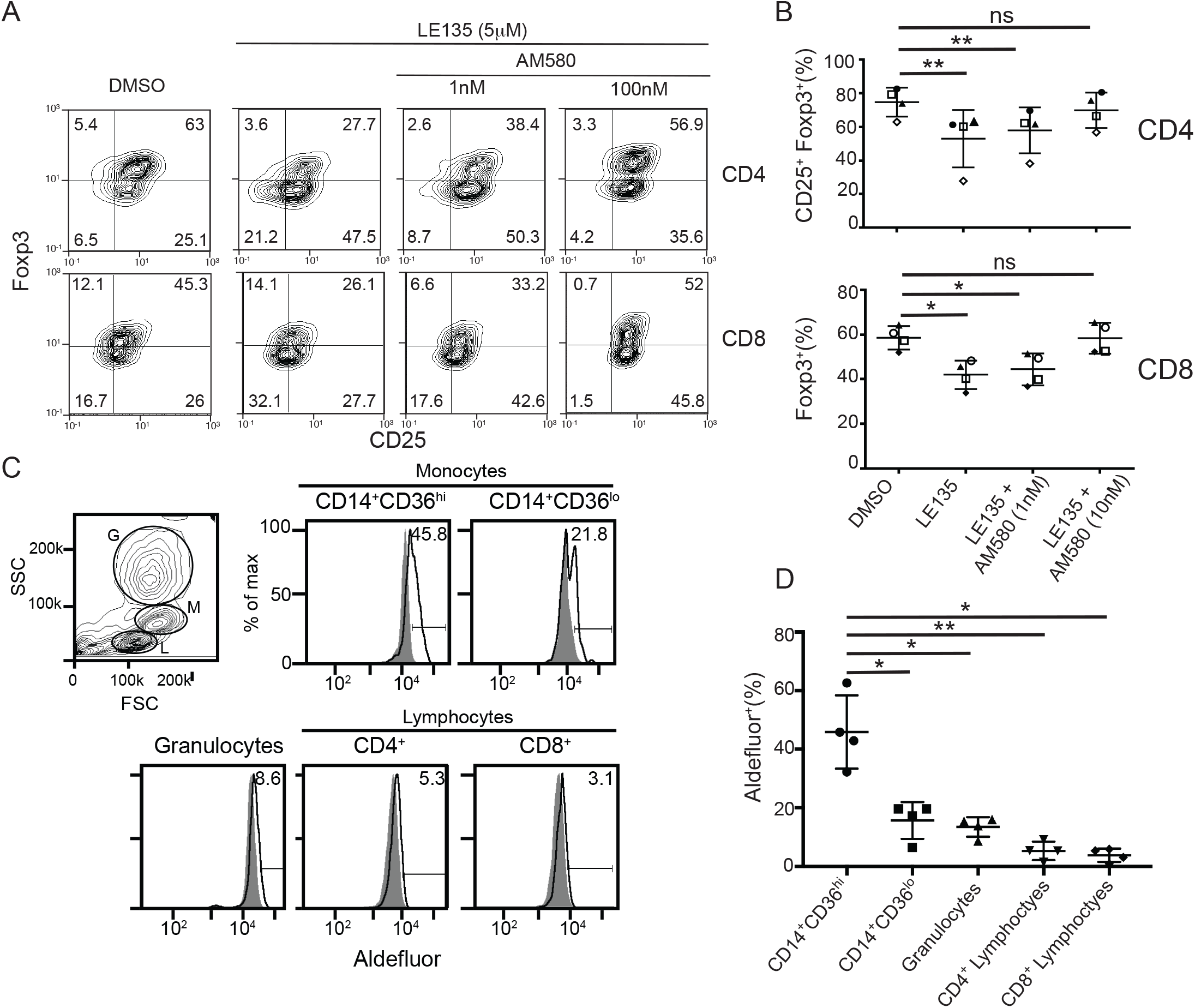
The role of retinoic acid in Foxp3^+^ cTregs generation. **(A-B)** Inhibition of Foxp3^+^ cTregs differentiation by a RAR antagonist. Total UCB cells were stimulated with anti-CD3, IL-2 and the RAR antagonist, LE135. Where indicated, the RAR agonist, AM580, was added in addition to LE135. **(A)** Representative plot of Foxp3 expression analyzed on CD4^+^ (upper panel) and CD8^+^ (lower panel) cells. **(B)** Summarized data, each shape represents a different donor; one-way ANOVA with Dunnett’s multiple comparison’s test, n=4. **(C)** Active ALDH expression by UCB monocytes. UCB cells were treated with the ALDEFLUOR™ assay and granulocytes (G), monocytes (M), and lymphocytes (L) were separately analyzed for ALDH activity. Histograms represent cells treated with the ALDEFLUOR™ reagent alone (open histogram) or in combination with a chemical inhibitor of ALDH as a measure of background fluorescence (solid histogram). **(D)** Summarized data from (C); one-way ANOVA with Dunnett’s multiple comparison’s test, n=4.

Retinoic acid is generated by serial oxidization of vitamin A by an alcohol dehydrogenase and aldehyde dehydrogenase (ALDH). Since UCB Foxp3^+^ T cell differentiation can be induced by purified CD14^+^CD36^hi^ monocytes, we hypothesized these monocytes express ALDH to generate retinoic acid. To test this, we determined the ALDH activity using a diffusible fluorescent dye that is retained by cells when oxidized by ALDH (ALDEFLUOR assay). We found a large percentage of CD14^+^CD36^hi^ and a small fraction of CD14^+^CD36^lo^ monocytes possess the ALDH activity (Fig. 5C, D). In contrast, only a minor fraction of granulocytic cells or CD4^+^/CD8^+^ T cells showed the appreciable ALDH activity. Together, our data suggest CD14^+^CD36^hi^ monocytes are a primary producer of retinoic acid among UCB cells, contributing to their ability to induce Foxp3^+^ T cell generation.

### 3.5 Generation of Foxp3^+^ T cells from naïve T cells by adult monocytes

Like UCB cells, adult PBMCs contain CD14^+^ CD36^hi^ monocytes, though at a lower rate (statistically not significant) (Fig. 6A). However, we observe much-reduced levels of Foxp3^+^ T cell development from total adult PBMCs (Fig. 1A). To decipher the mechanism by which adult immune system promotes more effector T cell responses, we first tested if naïve T cells-monocytes interactions in adult blood could lead to the generation of less Foxp3^+^ T cells compared to UCB. We examined if adult CD14^+^CD36^hi^ monocytes induce Foxp3^+^ T cells from cord blood at the comparable level to cord blood-derived monocytes (Fig. 6B). When we mix adult monocytes and UCB derived T cells and stimulate T cells with anti-CD3 Ab and IL-2 by the same method for UCB culture described above, we observe over 70% of cord blood T cells become Foxp3^+^ (Fig. 6B). The level of Foxp3^+^ T cells was comparable between cord blood-derived monocytes and adult blood-derived monocytes. Thus, the difference we observed in total PBMC in Foxp3^+^ T cell development is not due to loss of function by monocytes.

**Fig. 6.**
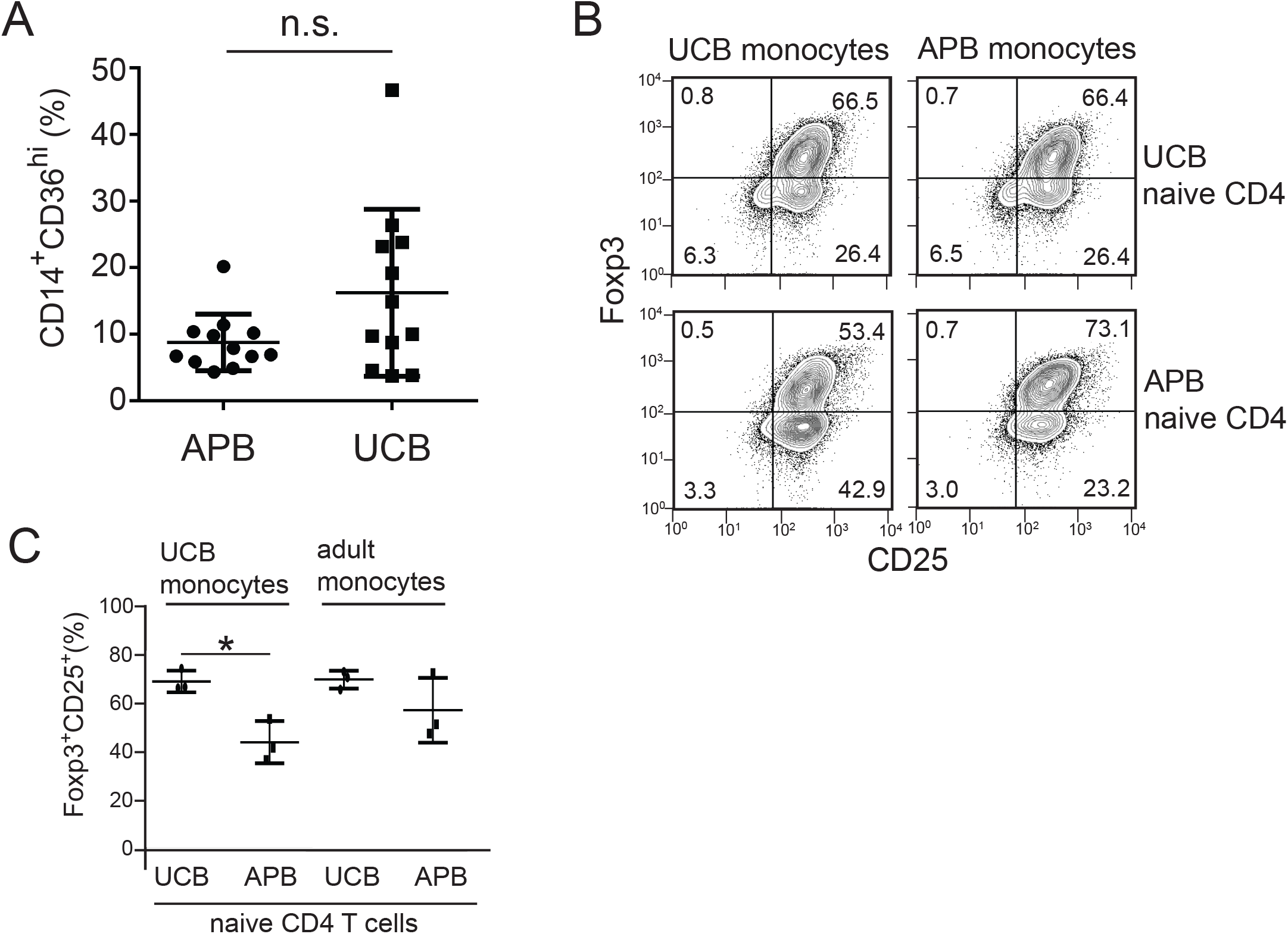
Comparison of adult and cord blood derived monocytes and T cells for cTreg induction. (**A**) Frequency of CD14^+^CD36^hi^ monocytes from freshly isolated umbilical cord blood (UCB) and adult peripheral blood (APB). Total nucleated cells from UCB or APB were analyzed for the expression of CD14 and CD36^hi^ phenotype. Percentage for each blood group is shown, n=12; n.s.=not significant. (**B**) Induction of cTregs from APB monocytes and APB naïve CD4 T cells. CD14^+^CD36^hi^ monocytes and naïve CD4^+^ T cells were purified from UCB or APB. Naïve CD4 T cells from UCB (upper panels) or APB (lower panels) were stimulated by anti-CD3 plus IL-2 in the presence of UCB monocytes (left column) or APB monocytes (right columns). After 2 weeks of culture, expression of Foxp3 and CD25 by CD4^+^ T cells were analyzed. A representative set of data from 3 samples. (**C**) The frequencies of Foxp3^+^CD25^+^ T cells developed form co-culture of UCB monocytes (left half) or APB monocytes (right half) with UCB or APB naïve CD4 T cells. (n=3). * p<0.05.

Next, we tested if adult naïve CD4 T cells have the same ability to become Foxp3^+^ T cells as compared to UCB naïve CD4 T cells since we observed some intrinsic differences between the UCB and adult blood naïve CD4 T cells previously [31]. When we stimulate adult naïve CD4 T cells by UCB monocytes, Foxp3^+^ T cell development was mildly lower than UCB derived monocytes (p<0.05) (Fig. 6C). We also observed a mild decrease (not statistically significant) in Foxp3^+^ T cell development when adult CD4 T cells were cultured with adult monocytes compared to UCB CD4 T cells. Together the data suggest that adult and UCB derived CD14^+^ monocytes are equally capable of inducing Foxp3^+^ Tregs, but naïve CD4^+^ T cells from adult PBMCs is less efficient to be Foxp3^+^ T cells than UCB naive CD4^+^ T cells.

## 4. Discussion

Our data demonstrate that antigen receptor stimulation of UCB T cells leads to predominant development of Foxp3 expressing CD4^+^ and CD8^+^ T cells (cTregs) with T cell suppressive ability. We found that Foxp3 is stably expressed over 60 days *in vitro*, which is substantially longer than tTregs. These cTregs express a distinctive set of surface antigens (CD26, CD31, OX40) when compared to APB CD4^+^CD25^+^Foxp3^+^ Tregs.

A unique feature of UCB derived Foxp3^+^ T cells is their surface antigen expression. While CD26 is expressed by adult T cells after activation [32], Foxp3^+^ cTregs cells maintain CD26 expression without continued antigen receptor stimulation. CD26, also known as dipeptidyl peptidase 4 (DPPIV), cleaves multiple ligands that contain proline or alanine residues at the N-terminus including chemokines and cytokines [33]. CD26 inactivates a group of insulinotropic proteins such as glucagon like peptide 1 (GLP1) [34] and negatively regulates glucose metabolism. CD26 enzymatic inhibitors are used clinically for treatment of insulin resistant diabetic patients [35]. Decreased glucose availability causes preferential expansion of Tregs [36]. Thus, a high level of CD26 expression may support expansion of cTregs in the fetal/neonatal environment.

In addition to its peptidase activity, CD26 binds adenosine deaminase (ADA) [37]. Adenosine is present at high level in fetal/neonatal blood and ADA deficiency causes severe immunodeficiency [38]. Expression of CD26 renders T cells resistant to suppressive effect of adenosine [39]. Therefore, CD26 expressed by UCB T cells and cTregs may play pivotal roles to maintain their reactivity *in vivo* by ADA-mediated removal of adenosine. It should be noted that high levels of constitutive CD26 expression are reported for Th17 cells [15] and mucosal associated invariant T cells (MAIT cells) [16, 40]. Enzymatic and genetic inhibition of CD26 reduced allograft rejection and IL-17^+^ cell development [41]. These data suggest that CD26 plays a role in maintenance/augmentation of a subset of T cell activation. While the functional relevance of CD26 expression by cTregs is unclear, cTregs may share some metabolic, activation, and/or epigenetic characteristics with Th17 and MAIT cells.

While CD26 appears to play T cell activating functions, loss of CD26 caused exacerbation of experimental autoimmune encephalomyelitis (EAE) [42]. In this model, production of Th1 type cytokines (IFN-γ and TNF) was highly elevated and the plasma level of TGF-β1 was significantly reduced. In human, prevalence of Crohn’s disease and Hashimoto’s thyroiditis among patients taking CD26 enzymatic inhibitor was significantly elevated compared to the control group, suggesting that CD26 plays a role in reduction/inhibition of autoimmune disorders [43].

In addition to CD26, CD8^+^ cTregs cells express CD31 (PECAM-1). CD31 is an inhibitory surface protein and contains an immunoreceptor tyrosine-based inhibitory motif (ITIM) in its cytoplasmic domain. CD31 plays a pivotal role in TGF-β mediated suppression of T cell activity such as inhibition of IFN-γ and Granzyme B expression [44]. CD31 is also a marker for recent thymic emigrants [45]. Loss of CD31 exacerbates inflammation in EAE, as well as collagen-induced arthritis (CIA) [46]. Expression of CD31 by practically all CD8^+^ and about 40% of CD4^+^ cTregs indicate that these T cells may be highly sensitive to TGF-β. In addition, CD31^+^ binds αvβ3 integrin and enhances leukocyte adhesion to endothelial cells, suggesting that CD31^+^ cTregs may be highly effective in tissue migration via endothelial adhesion. [47].

Expression of OX40 by a subset of cTregs raises several interesting possibilities. OX40 is not required for the development of tTregs [48], and ligation of OX40 reduced the suppressive functions of tTregs [49]. Moreover, the signal from OX40 inhibits Foxp3 expression by activated naive CD4^+^ T cells in the presence of TGF-β and blocked iTreg differentiation [50] Together, OX40 signaling inhibits regulatory T cells’ function and development. Constitutive expression of OX40 by a subset of cTregs indicates these cTreg’s function may be controlled negatively by antigen presenting cells expressing OX40 ligand (CD252) such as monocyte-derived dendritic cells [51] and activated B cells [52]. The data also suggest that agonist of OX40 could be used to reduce cTregs functions and potentiate immune responses in the neonate.

The different surface antigens on cTregs and adult Tregs may be used as markers to distinguish the two populations in settings where adult and fetal cells mix, such as during pregnancy and neonate [53]. This is especially true of CD26, due to its expression by virtually all UCB-derived CD4^+^ and CD8^+^ cTregs, but not APB Tregs.

Of note, it has not been previously described prior to this study that monocytes have a role in the induction of stable Tregs. However, our data demonstrate that for UCB T cells to become Foxp3^+^ T cells, CD14^+^CD36^hi^ monocytes play an essential role. Mechanistically, we found that these monocytes provide TGF-β and retinoic acid to the UCB T cells. Depletion of CD14^+^CD36^hi^ monocytes from UCB abrogated cTreg differentiation and activation of Smad2/Smad3. We also observed elevated ALDH activities in CD14^+^CD36^hi^ monocytes. The data suggest that CD14^+^CD36^hi^ monocytes are potent immuno-modulatory antigen presenting cells. Importantly, we found that induction of Foxp3^+^ Tregs is not a unique feature to UCB derived monocytes. While naïve CD4 T cells from APB are less prone to be Foxp3^+^ T cells than UCB naïve CD4 T cells, adult peripheral blood monocytes are equally capable of inducing Foxp3^+^ Tregs.

In addition to CD4^+^ cells, CD14^+^CD36^hi^ monocytes induce CD8^+^ T cells to become Foxp3^+^ Tregs. CD8^+^ T cells with suppressive functions have been recognized for many years. Indeed, suppressor T cells were initially identified in the CD8^+^ subset [54-56]. Recently, many reports identified CD8^+^ regulatory T cells involved in normal and pathogenic immune responses [57, 58]. Although the human thymus and peripheral blood contains functional CD8^+^ Tregs [59], the conditions for CD8^+^ Treg induction has not been well established. The data suggest that monocytes from adult blood can be a useful tool for induction of regulatory T cells to treat autoimmune disorders. Conversely, inhibition of monocytes-mediated Treg differentiation could help immunotherapy against cancers.

Monocytes are capable of surveying the peripheral vascular environment and transporting antigen back to distant organs [25, 60, 61]. It is largely accepted that after stimulation and extravasation from the bloodstream, monocytes differentiate into macrophages or dendritic cells and are no longer monocytes, but “monocyte-derived” cells [61]. Because monocytes are circulating cells, antigen encountered prior to extravasation can be retained, transported and presented in other organs [61]. *In vivo* studies on CD14^+^CD36^hi^ monocytes will potentially provide more details on their functions in the future.

As demonstrated by landmark works by Owen, Medawar, and their colleagues [1] [2], the fetal immune system is programmed to generate immunological tolerance. Development of cTregs maybe one of the mechanisms that establish long-lasting tolerance against foreign antigens. The tolerogenic nature of perinatal immune system may be an essential process for newborn babies to establish symbiotic conditions with commensal microorganisms. While this is generally beneficial for the infant, the tolerogenic nature of perinatal immune system plays a double sided-sword for the health of a newborn. The recent increase in fetal vertical infections by Zika virus clearly depicts the risk of this process. Like other TORCH infections (Toxoplasma, Other, Rubella, Cytomegalovirus, Herpes virus) that are transmitted during gestation, Zika can permanently devastate the affected child. To protect fetus from these pathogenic microorganisms, elucidation of the fetal immune responses against pathogens is urgently needed. Further analysis of Foxp3^+^ cTregs from the fetus and neonate will help developing better prevention/treatment for neonatal infections.

## Acknowledgement

The authors thank Patricia Simms (Loyola FACS core facility) for cell sorting. The authors would also like to thank the nurses at Gottlieb Memorial Hospital and Loyola University Medical Center for their critical participation in collection of UCB. This work was supported by NIH R01 AI055022 (M.I), AI I100129 (M.I. and K.L.K), The Uehara Memorial Foundation research fellowship (Y.S.), NIH T32 AA013034 (K.E.J.), and the Van Kampen Cardiopulmonary Research Fund (M.I.).

## Supplementary figures

**sFig. 1.**
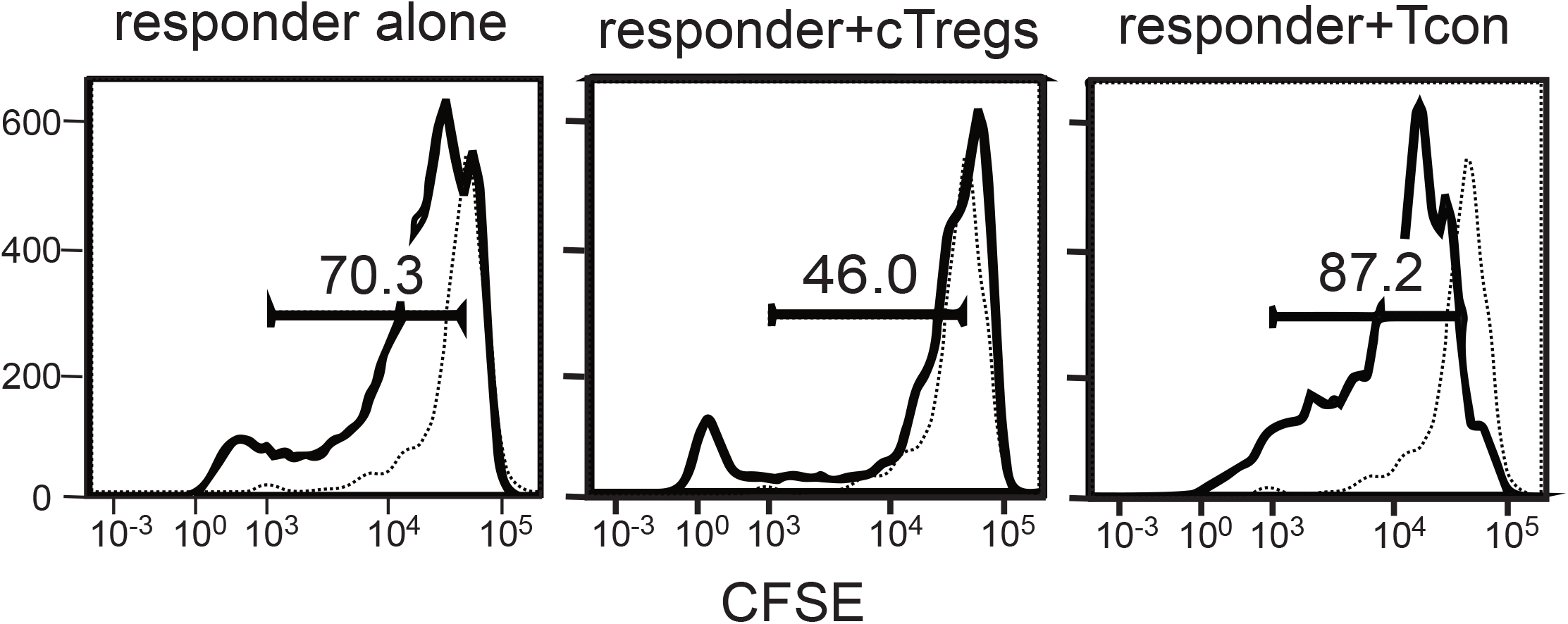
*In vitro* suppression of T cell proliferation by CD4^+^ cTregs or conventional T cells. UCB CD4^+^ T cells were stimulated by syngeneic APCs in the absence (left panel) or presence of cTregs (middle panel), CD4^+^CD25^-^ conventional T cells (right panel). Conventional T cells were prepared by activation of CD4^+^CD25^-^ T cells by anti-CD3 and anti-CD28 antibody coated beads (5 days). Proliferation of responder T cells were measured by the dilution of CFSE. Dotted lines show the response of unstimulated cells. Data representative of 3 independent experiments.

**sFig. 2.**
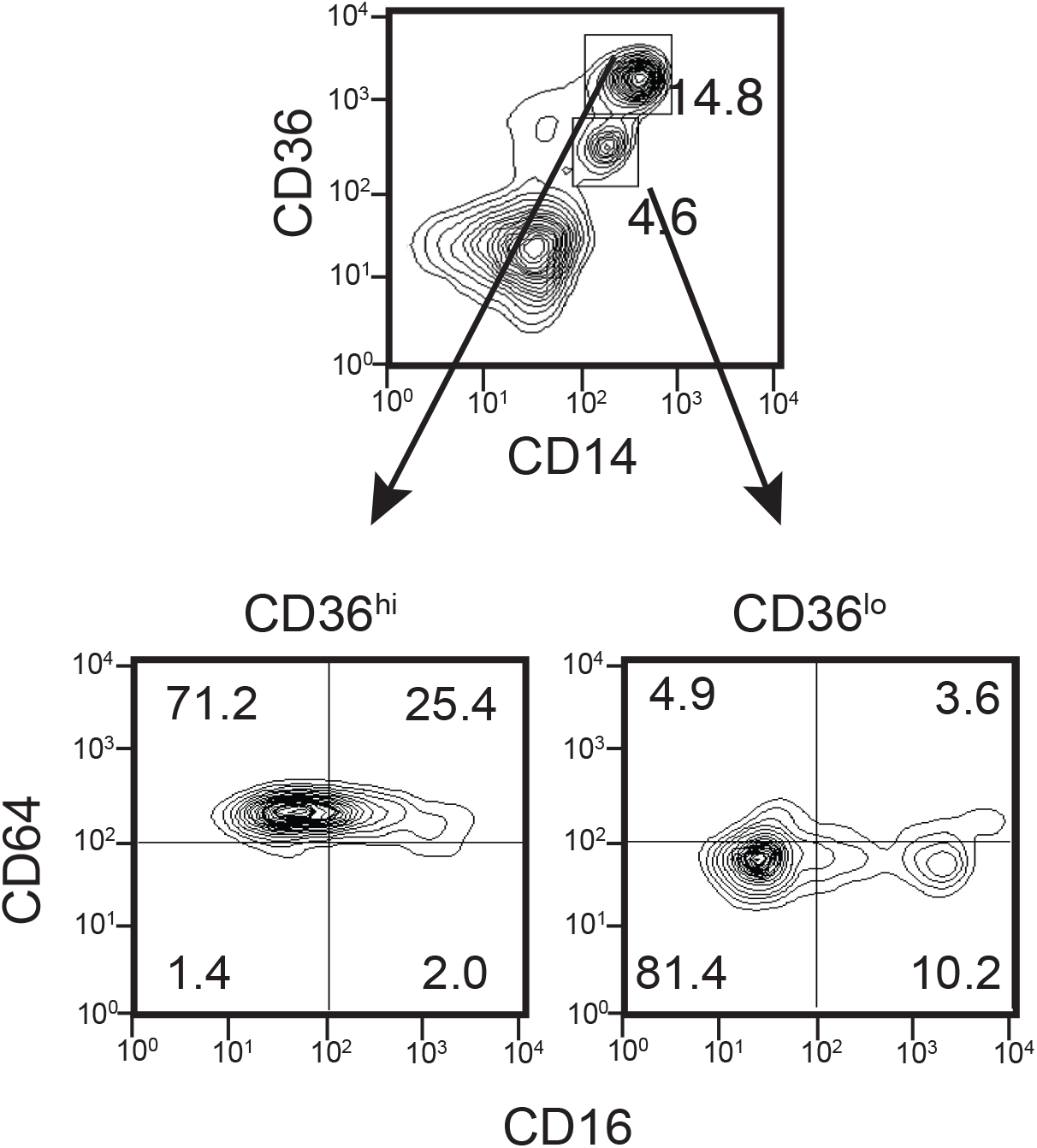
Analysis of surface CD16 and CD64 on freshly isolated UCB CD14^+^CD36^hi^ and CD14^+^CD36^lo^ monocytes. Data are representative of at least three independent experiments.

**sFig. 3.**
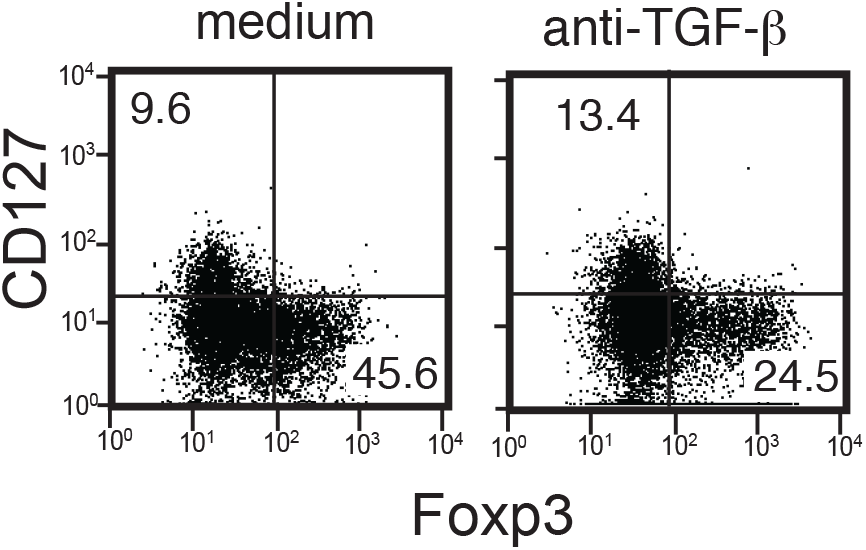
Inhibition of Foxp3^+^ CD127-T cell development by anti-TGF-β antibody. Total UCB T cells were stimulated by anti-CD3 and IL-2 in the absence (left panel) or presence (right panel) of an anti-TGF-β blocking Ab, then cultured for 14 days as described in Fig. 1. Representative data from 3 samples.

**sFig. 4.**
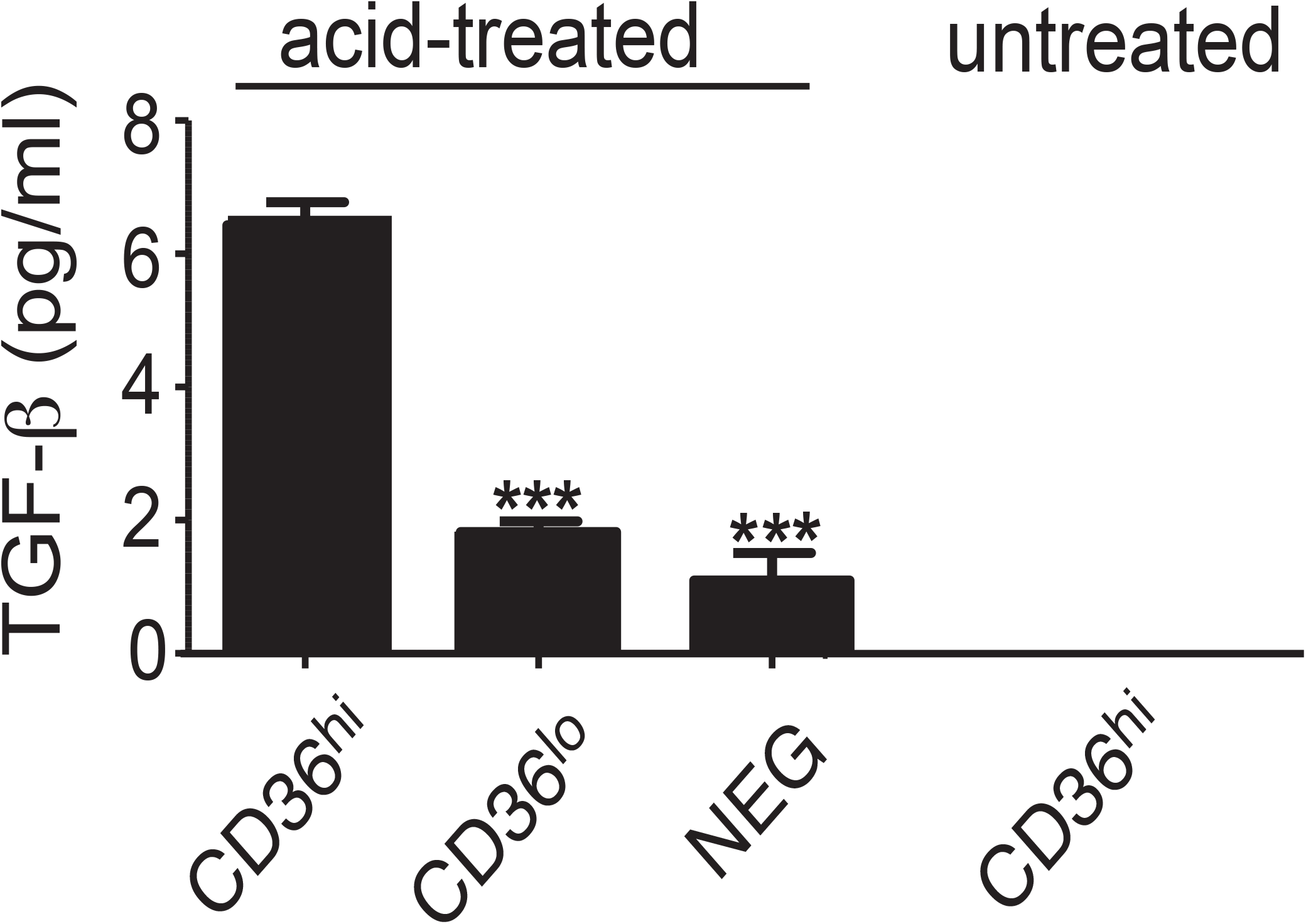
Production of soluble TGF-β by CD14^+^CD36^hi^, CD14^+^CD36^lo^, or CD14-UCB cells. Each cell subset was cultured for 24 hours and the culture supernatant analyzed for TGF-β using a reporter-based bioassay system [11]. The level of total (acid-treated) and active (untreated) TGF-β was determined. Active TGF-β was below the detectable limit of the assay. An asterisk indicates a significant decrease (p<0.001) in total TGF-β production compared to CD14^+^CD36^hi^ cells by one-way ANOVA with Tukey’s post test, n=6.

